# Complex patterns of biological connectivity highlight risks of local depletion in an Australian fishery

**DOI:** 10.64898/2026.07.15.738810

**Authors:** Lauren Brown, Nick S. Whiterod, Justin Rizzari, Thomas C. Barnes, John R. Morrongiello, Jason Lieschke, Adam D. Miller

## Abstract

Sustainable management of commercial and recreational fisheries depends on accurately resolving population connectivity, across both ecological and evolutionary timescales. However, dispersal can vary markedly among life stages, making stock connectivity difficult to resolve using single-method approaches that often differ in spatio-temporal resolution. Here, we integrated population genomics, otolith stable isotope chemistry, and mark–recapture analyses to provide a multi-faceted assessment of stock connectivity in mulloway (*Argyrosomus japonicus*). Mulloway are a commercially, culturally, and recreationally important estuary associated fish distributed throughout the Indo-Pacific region, including south-eastern Australia where this study was conducted. Genome-wide single nucleotide polymorphism (SNP) analyses revealed significant genetic differentiation between regions influenced by different current systems, but limited structure within regions across distances exceeding 900 km. In contrast, otolith δ^13^C and δ^18^O signatures revealed fine-scale spatial structuring among estuaries, consistent with prolonged occupancy of local habitats. Mark–recapture analyses supported this interpretation, with most fish exhibiting strong estuarine fidelity over extended periods despite occasional long-distance coastal movements. Reconstructed age structures from fish otoliths revealed remarkably similar cohort composition among estuaries, with populations dominated by cohorts originating from a major recruitment pulse centred on 2011–2012, likely associated with a broad-scale flood-driven spawning and recruitment event. Together, our findings indicate that mulloway fisheries function as regionally connected networks of partially independent estuarine assemblages, where strong local residency is periodically offset by dispersive individuals and episodic recruitment events that maintain long-term demographic and genetic connectivity. Consequently, local estuarine populations may be vulnerable to localised depletion despite broader regional connectivity, particularly where sustained fishing pressure coincides with reductions in freshwater flows that constrain spawning and recruitment. More broadly, our study demonstrates the value of integrating complementary approaches to identify biological connections and define meaningful management units in species with complex life histories.

## 1 INTRODUCTION

Sustainable fisheries management depends on accurately defining biological stock structure to assess resilience to harvest pressure and environmental disturbance (Fao 2023; Ices 2025). Here, delineating population boundaries and identifying associated recruitment dynamics is essential for implementing spatially explicit management that reduces risks of local and regional depletion, identifies critical habitats for protection, and guides recovery interventions such as restocking (Reiss *et al*. 2009; Kerr *et al*. 2017). However, connectivity patterns are rarely static, and often complex. Dispersal capacity and spatial patterns of habitat use can vary markedly across life stages, with larvae, juveniles, sub-adults and adults often occupying different habitats and exhibiting contrasting movement behaviours. Atlantic cod (*Gadus morhua*) exemplify this, with pelagic larvae dispersing widely, juveniles using shallow coastal nurseries, and adults undertaking seasonal spawning migrations while also often displaying strong site fidelity (Zemeckis *et al*. 2014a; André *et al*. 2016; Wright *et al*. 2021). The strength and directionality of an individual’s movement and spatial range is therefore strongly life-stage dependent, complicating stock delineation and management in Northwest Atlantic cod fisheries (Kerr *et al*. 2014; Zemeckis *et al*. 2014b). A reliance on assessment methods that fail to capture these connectivity dynamics risks mischaracterising stock structure, undermining management effectiveness, and increasing the likelihood of serial depletion.

A range of tools are used to estimate biological connections in exploited fish species. Population genetic approaches characterise spatial patterns of gene flow (Waples *et al*. 2008; Benestan 2020), otolith elemental and isotopic analyses provides insight into habitat use, diet and natal origins (Catalán *et al*. 2018; Wright *et al*. 2018), and tagging or telemetry enables direct observation of animal movement in both time and space (Crossin *et al*. 2017; Brownscombe *et al*. 2019). Each method captures distinct temporal and biological processes: genetic data reflect intergenerational connectivity but may obscure contemporary demographic independence; otolith signatures reveal habitat use but not reproductive exchange; and tagging studies are often limited in replication and biased toward larger individuals and short timeframes (Cadrin *et al*. 2013; Sarakinis *et al*. 2024). Integrating these complementary tools is therefore essential for resolving complex stock dynamics. A well-known example is Pacific salmon (*Oncorhynchus* spp.) management in the North Pacific, where integrating genetics, otolith microchemistry and large-scale tagging has resolved stock structure and migration pathways by assigning natal origin, reconstructing early marine history, and estimating migration, survival and harvest patterns (Brennan *et al*. 2015; Carbonneau *et al*. 2024; Goertler *et al*. 2024). However, such integrated assessments remain uncommon, as these methods are often applied in isolation, raising concerns about potential fisheries mismanagement.

Mulloway (*Argyrosomus japonicus*) is a large coastal sciaenid fish species with a broad distribution extending throughout southern Australia, the north-western Pacific Ocean and the Indian Ocean (Griffiths & Heemstra 1995; Silberschneider & Gray 2008). In Australia, the species occurs across more than 6000 km of the southern coastline extending from Western Australia to Queensland, where it is common to estuaries and the lower reaches of rivers, as well as inshore rocky reef and ocean beach habitats (Kailola *et al*. 1993; Taylor *et al*. 2006a; Silberschneider & Gray 2008). The species has a long history of exploitation in Australia, from Indigenous harvest through to modern fishery expansion, decline, and reform. Historically targeted in estuaries and nearshore coastal waters, the fishery intensified during the 1980s and 1990s, particularly in New South Wales and South Australia, with commercial landings peaking during this period (Silberschneider & Gray 2008; Hughes 2023). Growing fishing pressure, combined with habitat degradation and altered freshwater flows that led to variable recruitment success, contributed to declining mulloway fishery productivity from the early 2000s onward across much of its Australian range. Commercial catches subsequently fell well below historical highs, and in some regions, mulloway were classified as “overfished” or “depleted” (Silberschneider & Gray 2008; Stewart *et al*. 2015; Earl *et al*. 2018). Similar trends have occurred in South Africa, where sustained fishing pressure and uncontrolled effort led to collapse of the stock, now estimated to be less than 5% of virgin spawning biomass (Griffiths 1997; Winker *et al*. 2015; Mirimin *et al*. 2016). In response, Australian management agencies introduced tighter size and bag limits, seasonal closures, spatial restrictions (DPIRD 2025), and effort controls, alongside strategic restocking programs in several states (Taylor *et al*. 2006b; Recfishwest 2024; Becker *et al*. 2026). The trajectory of mulloway fisheries highlights the risk of localised depletion and reinforces the importance of resolving stock structure and connectivity to guide spatially explicit, evidence-based management.

Across their range, mulloway exhibits a complex life history that complicates stock assessment and management (Griffiths 1997; Ferguson *et al*. 2008; Ferguson *et al*. 2011; Mirimin *et al*. 2016; Knight *et al*. 2025). In Australia, spawning occurs in nearshore coastal waters often adjacent to large river mouths between November and March, typically following major rainfall or freshwater flow events that provide low-salinity cues for adult aggregations (Ferguson *et al*. 2008; Silberschneider *et al*. 2009; Taylor *et al*. 2014). These infrequent, flow-driven spawning events likely play a disproportionate role in sustaining regional connectivity, genetic diversity, and replenishment of estuarine stocks, while also contributing to high recruitment variability (Ferguson *et al*. 2008; Taylor *et al*. 2014; NSW Department of Primary Industries 2016). Pelagic eggs and newly hatched larvae are subsequently dispersed by coastal currents, potentially facilitating long-distance connectivity (Silberschneider & Gray 2008; Silberschneider *et al*. 2009). Consistent with this, population genetic studies in Australia report limited genetic structure within regions linked by common current systems, but differentiation between regions influenced by different currents (Archangi 2008; Barnes *et al*. 2016). In contrast, movement becomes more restricted following recruitment into estuarine nursery habitats. Otolith microchemistry and acoustic telemetry studies indicate juveniles are largely sedentary with small home ranges, while many sub-adults and adults display strong estuarine fidelity despite occasional movements exceeding 1000 km (Taylor *et al*. 2006b; Ferguson *et al*. 2011; Lieschke 2019; Russell *et al*. 2021; Hughes *et al*. 2022). This ontogenetic shift in connectivity suggests that broad-scale genetic homogeneity may mask fine-scale demographic independence among estuaries. Such dynamics increase the risk of localised or serial depletion between episodic recruitment events, particularly where declines in freshwater flows due to river regulation or climate change disrupt spawning cues and nursery habitat quality (Taylor *et al*. 2014).

Although population genetics, otolith chemistry and movement tracking have each provided valuable insights into mulloway spatial ecology, these approaches have largely been applied in isolation and often in different regions or time periods. As a result, broad-scale genetic connectivity and fine-scale movement studies have rarely been evaluated simultaneously within the same estuarine systems, creating uncertainty about how intergenerational gene flow aligns with contemporary demographic exchange. This disconnect has limited our ability to determine whether genetically homogeneous stocks function as demographically independent units at management-relevant scales. Here, we integrate population genomics, mark–recapture data, and otolith stable isotope analyses to resolve connectivity among mulloway stocks from south-eastern Australia. Specifically, we test whether broad-scale genetic homogeneity masks fine-scale demographic independence, quantify movement and habitat use among estuaries, and assess implications for stock resilience. By combining intergenerational and contemporary connectivity measures within a common geographic framework, this study provides a robust, multi-method approach to inform spatially explicit management of mulloway fisheries and underscores the importance of integrating complementary tools when stock dynamics are complex and life-stage dependent.

## 2 METHODS

This study integrated three complementary datasets, population genomic, otolith stable isotope, and mark–recapture data, to assess stock connectivity and movement dynamics of mulloway (*A. japonicus*) across south-eastern Australia. The majority of data used were generated through a community-based citizen science program led by Nature Glenelg Trust between 2014 and 2020, funded by the Victorian Fisheries Authority through recreational fishing licence grants (Project 15/15278). The program engaged recreational anglers across south-eastern South Australia and Victoria in long-term tagging and biological sampling to improve understanding of stock structure, movement, growth, and recruitment dynamics. More than 150 anglers contributed biological samples and donated fish frames, while trained volunteers undertook tagging across multiple estuaries and coastal waters, generating valuable data on estuarine residency, long-distance movement, growth, and recapture patterns. Beyond its scientific value, the program fostered strong community engagement by directly involving recreational fishers in fisheries research and management.

### 2.1 Population genomics

#### 2.1.2 Biological sampling

Tissue biopsies (fin clips) were collected for genomic analysis from 6–33 individual mulloway from 14 estuaries in south-eastern Australia (Figure 1; Table 1). Samples from South Australia and New South Wales was collected by recreational anglers and commercial operators between 2009 and 2012 (Barnes *et al*. 2016). Most Victorian samples were collected between 2014 and 2016 by recreational anglers participating in the citizen science project funded by the Victorian Fisheries Authority, with supplementary sampling conducted by the Arthur Rylah Institute in 2016 (Table 1). Individual tissue biopsies were obtained using flame-sterilised scalpels and forceps, preserved in 2 mL screw-cap vials containing 100% ethanol, and stored at room temperature until genetic analysis.

**Figure 1.**
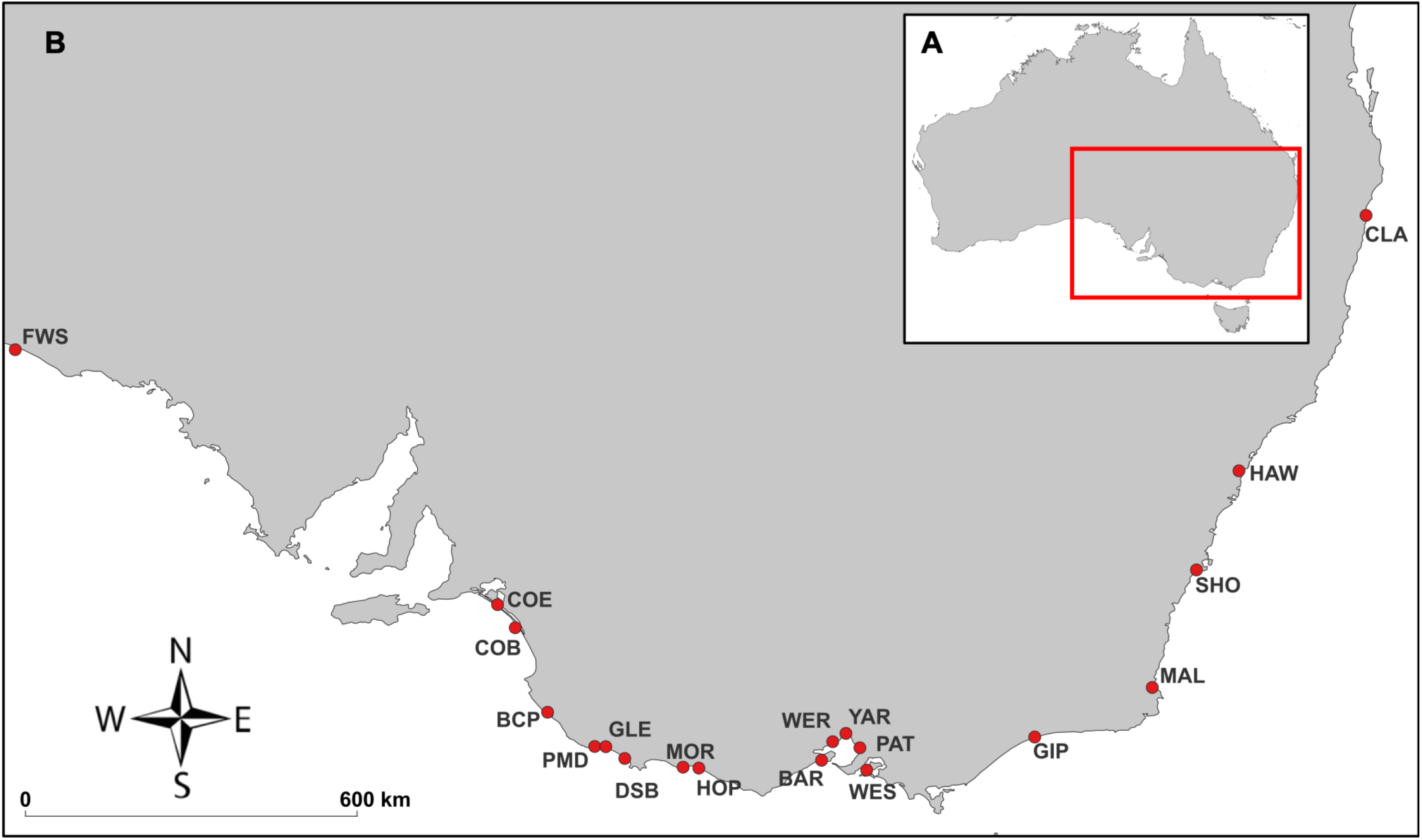
Sampling locations of mulloway (*Argyrosomus japonicus*) across south-eastern Australia used in population genomic, otolith stable isotope, and movement analyses. Sampling sites are shown for estuarine and coastal waterways spanning South Australia, Victoria, and New South Wales. Site abbreviations correspond to waterways detailed in Table 1, which also summarises sample sizes and data availability for genomic (n Genetics), otolith stable isotope (n Isotopes), and movement datasets, including tagged individuals (n Marked) and recaptures (n Recaptures).

**Table 1.** Sample sizes and data availability for mulloway (*Argyrosomus japonicus*) collected across southern and eastern Australian waterways, including individuals used for population genomic (*n* Genetics), otolith stable isotope (*n* Isotopes), movement (*n* Movement; tagged fish and recaptures in parentheses), and demographic (*n* Demographics; age structure, cohort composition, and age–length) analyses.

| Location | Code | GPS Coordinates | *n*<br>Genetics | *n*<br>Isotopes | *n*<br>Movement | *n*<br>Demographics |
| --- | --- | --- | --- | --- | --- | --- |
| Far West SA | FWS | -31.600, 131.397 | 30 | 0 | 0 (0) | 0 |
| Coorong Beach | COB | -36.120, 139.523 | 33 | 9 | 25 (4) | 9 |
| Coorong Estuary | COE | -35.745, 139.236 | 6 | 3 | 8 (1) | 0 |
| Beachport | BCP | -37.494, 140.049 | 0 | 0 | 19 (3) | 0 |
| Port MacDonnell | PMD | -38.051,<br>140.815 | 0 | 0 | 59 (25) | 0 |
| Glenelg River | GLE | -38.054, 140.995 | 31 | 56 | 581 (104) | 363 |
| Discovery Bay | DSB | -38.246,<br>141.303 | 0 | 0 | 15 (2) | 4 |
| Hopkins River | HOP | -38.395, 142.246 | 32 | 55 | 40 (5) | 78 |
| Moyne River | MOR | -38.387,<br>140.995 | 0 | 0 | 13 (3) | 13 |
| Barwon River | BAR | -38.275, 144.501 | 33 | 54 | 18 (1) | 156 |
| Werribee River | WER | -37.973, 144.682 | 31 | 0 | 0 (0) | 0 |
| Yarra River | YAR | -37.838, 144.897 | 6 | 9 | 25 (4) | 9 |
| Patterson River | PAT | -38.072, 145.124 | 30 | 39 | 81 (21) | 39 |
| Westernport Bay | WES | -38.434, 145.234 | 10 | 5 | 0 | 13 |
| Gippsland Lakes | GIP | -37.897,<br>147.964 | 0 | 2 | 0 | 0 |
| Mallacoota | MAL | -37.091, 149.874 | 10 | 3 | 0 | 8 |
| Shoalhaven River | SHO | -35.180, 150.592 | 24 | 0 | 0 | 0 |
| Hawkesbury River | HAW | -33.569, 151.282 | 24 | 0 | 0 | 0 |
| Clarence River | CLA | -29.417, 153.347 | 32 | 0 | 0 | 0 |

#### 2.1.2 DNA Extraction and SNP Genotyping

A total of 332 tissue biopsies representing 14 locations were used for genomic analysis (refer to Table 1 for site and source details, and Figure 1). Genomic DNA was extracted from 10 mg of muscle tissue with DNA Blood and Tissue kits (QIAGEN, Venlo, Limburg, Netherlands), and reduced representation genome libraries were prepared with a modified genotyping by sequencing (GBS) protocol of Elshire *et al*. (2011). Five hundred nanograms of genomic DNA from each individual was digested in a 20 μL reaction containing four units of SbfI and MseI for 2 h at 37 °C, without a heat kill step. Digestion products were then ligated to modified P1 and P2 adapters with 80 unique barcode combinations to allow for subsequent multiplexing of all individuals. Fifty microlitre ligations were performed containing the SbfI and MseI digested DNA, 1.125 ng of P1 and P2 adapters, 400 units of T4 ligase and 19 units of T4 buffer (New England Biolabs, Beverly MA, USA). Ligations were incubated at 16 °C for 90 min followed by a 30 min of denaturation at 80 °C. Adapter ligated DNA fragments were purified using a Qiagen MinElute PCR purification kit (Redwood City, CA, USA), eluted in 20 μL of ddH20 and subsequently used for PCR amplification. Fifty microlitre PCRs were performed using 29 MyTaqTM HS Mix (Bioline, Taunton, MA, USA), and containing 0.2 μM each of Illumina Dual Index Sequencing Primers 1 & 2 (Illumina Inc., San Diego, CA, USA) and 10 μL of above purified DNA. PCR conditions were as follows: 95°C for 1 min, 24 cycles of 95 °C for 30 s, 65 °C for 30 s, 72 °C for 30 s and a final extension step of 72 °C for 5 min. DNA quantitation and qualitative analysis of individual PCR products were performed on a MCE - 202 MultiNA with a DNA-1000 kit (Shimadzu, Kyoto). The 332 individual libraries were pooled equimolar into four groups (pools of 42 samples) and sequenced on separate lanes of the Illumina HiSeq 4000 sequencing platform.

Raw sequences were first processed using the Trimmomatic V0.36 program (Bolger *et al*. 2014) by trimming the raw reads to 80 bp length and discarding all reads that had a Phred score below 20. We used the de novo program from STACKS 1.44 (Catchen *et al*. 2013) to create a catalogue of SNPs and genotypes for all individuals. Single nucleotide polymorphisms (SNPs) were subsequently called by retaining a single randomly selected SNP per tag, and only bi-allelic SNPs present in at least 80% of individuals in all populations with a minimum minor allele frequency of 0.05. Individuals or SNP loci with more than 20% missing data were excluded from further analysis.

#### 2.1.3 Tests for Population Genetic Structure

Population genomic analyses were conducted in R v4.2.1 (R Core Team 2022) using the packages *adegenet* (Jombart & Ahmed 2011; Jombart *et al*. 2018), *hierfstat* (Goudet 2005), *pegas* (Paradis 2010), *poppr* (Kamvar *et al*. 2014), *vegan* (Oksanen *et al*. 2022), *StAMPP* (Pembleton *et al*. 2013), and *ade4* (Dray & Dufour 2007). Genetic diversity statistics, including observed (*H*_O_) and expected heterozygosity (*H*_E_), and Weir and Cockerham’s inbreeding coefficient (*F*_IS_) were calculated using the function `basic.stats` in *hierfstat*.

Global estimates of population differentiation (*F*_ST_) and *F*_IS_, including associated 95% confidence intervals, were estimated using the Weir and Cockerham (1984) approach implemented in the function `wc`, while pairwise population differentiation was estimated using `pairwise.WCfst`. Patterns of population structure were further explored using discriminant analysis of principal components (DAPC) implemented in *adegenet*. The optimal number of genetic clusters was identified using the `find.clusters` function based on successive k-means clustering and Bayesian Information Criterion (BIC) minimization. DAPC was then performed using retained principal components to visualise patterns of genetic clustering and individual assignment probabilities among populations. Analysis of molecular variance (AMOVA) was conducted using `poppr.amova` in the *poppr* package to partition genetic variation among populations and DAPC-inferred regional clusters, with significance assessed using 999 permutations via `randtest`.

Population structure and recent shared ancestry among mulloway individuals were further investigated using fineSTRUCTURE (Raj *et al*. 2014). Analyses were conducted on the filtered SNP dataset using a Bayesian clustering framework for coancestry estimation among individuals. Clustering analyses were run for *K* = 1–10 using Markov chain Monte Carlo (MCMC) simulations with 100,000 burn-in iterations, 100,000 sampling iterations, and a thinning interval of 1,000 iterations. The optimal number of clusters was assessed by comparing model support across *K* values using the `chooseK.py` utility, which evaluates marginal likelihood and model complexity to identify the most parsimonious clustering solution. Hierarchical relationships among inferred clusters were subsequently reconstructed using the fineSTRUCTURE tree-building algorithm to generate coancestry dendrograms.

Finally, relative directional migration among populations was assessed using the `divMigrate` function implemented in the R package diveRsity (Keenan *et al*. 2013). Migration estimates were inferred from asymmetric genetic differentiation based on Nei’s *G*_ST_ statistic using all SNP markers. Analyses were conducted using the `gst` option with a filter threshold of 0.05 and 1,000 bootstrap replicates to estimate relative connectivity strength among sampled waterways. Resulting migration values were scaled between 0 and 1, where higher values indicated stronger inferred relative connectivity within the dataset. Because divMigrate infers directional connectivity indirectly from asymmetries in allele frequencies, migration estimates were interpreted qualitatively as indicators of relative gene flow and connectivity structure rather than direct demographic migration rates.

### 2.2 Otolith stable isotope analysis

#### 2.2.1 Otolith stable isotope analysis

Otoliths samples for stable isotope analysis were extracted from 5 to 55 individual adult fish frames between 2014 and 2016 from 8 locations, spanning South Australia and Victoria (Table 1). Frozen fish frames were provided by recreational anglers involved in the citizen science project described above. Sagittal otolith pairs were removed from individual fish using ceramic forceps, gently cleaned and dried, and stored in labelled envelopes for subsequent microchemistry analyses.

#### 2.2.2 Isotope measurements

Otolith stable isotope measurements were obtained from 235 fish collected across nine waterways spanning south-eastern South Australia and Victoria (Table 1). Sample sizes averaged 26.1 individuals per waterway and ranged from 5 individuals (Westernport Bay) to 55 individuals (Glenelg River and Hopkins River). From each fish, one sagitta was selected at random, cleaned and homogenized by being crushed to a powder in an agate mortar and pestle. Whole sagittae were deproteinated to remove proteins contained in the otolith carbonate matrix. These proteins could potentially degrade to form CO_2_ on analysis of the carbon and oxygen stable isotopes and therefore potentially interfere with the measurement of the carbon. Powdered sagittae were analysed for carbon (^13^C/^12^C) and oxygen (^18^O/^16^O) stable isotopes and reported using the international standard delta (δ) notation relative to the PDB-1 standard for carbonates (δ^13^C and δ^18^O). Measurements were conducted on a continuous-flow mass spectrometer (Analytical Precision AP2003) housed at the Stable Isotope Laboratory, School of Geography, University of Melbourne, Australia. Subsamples of 0.7 ± 0.1 mg were digested in 105% orthophosphoric acid at 70°C and measurements made on the evolved CO_2_. Sample results were converted to the VPDB scale using the known values of a Cararra Marble house standard (NEW1) previously calibrated to the VPDB scale using NBS19 and NBS18. A second house standard (NEW12), also calibrated against NBS18 and NBS19, was used as a cross-check in case scale correction was required (which was not required). The repeatability precision (1σ) on the AP2003 is ≤0.05‰ for δ^13^C and ≤0.10‰ for δ^18^O.

#### 2.2.3 Statistical analysis

To investigate spatial structuring in otolith isotope chemistry among mulloway capture waterways, both univariate and multivariate classification approaches were applied to otolith δ^13^C and δ^18^O isotope data. Statistical analyses were initially conducted on all 235 individuals, and subsequently on a subset of 149 individuals with complete age and total length data to assess ontogenetic effects on spatial structure. Finally, we analysed isotope data from 123 individuals with available sex information to explore any sex-specific effects. Ontogenetic stages were classified using age and total length (TL) criteria, with fish aged 2–4 years and 50–70 cm TL classified as sub-adults (n = 25), and fish >4 years and >70 cm TL classified as adults (n = 124). Sex-specific analyses comprised 75 females and 48 males. Sample sizes among ontogenetic stages and sexes were uneven, with several waterways represented by relatively small numbers of individuals within demographic groups. Consequently, ontogenetic and sex-specific analyses were considered exploratory and interpreted cautiously.

To initially assess spatial differences in isotope chemistry among waterways, one-way analyses of variance (ANOVA) were conducted separately for δ^13^C and δ^18^O isotope ratios using waterway as the grouping factor. Analyses were conducted in R v4.2.1 (R Core Team 2022) using the base *stats* package and implemented through RStudio v2023.6.1.524 (R Studio Team 2023).

Supervised classification analyses were then conducted using random forest algorithms implemented in the R package *randomForest* (Liaw & Wiener 2002) to assess the ability of otolith isotope signatures to correctly assign individuals to their known waterway of capture. Random forest classification was selected due to its ability to accommodate non-linear relationships among predictor variables and its robustness when analysing ecological datasets containing complex spatial structure (Breiman 2001; Cutler *et al*. 2007). Prior to analysis, δ^13^C and δ^18^O isotope variables were standardised to account for differences in variable scale. Random forest models were fitted using capture waterway as the response variable and isotope ratios as predictor variables. Model performance was evaluated using the out-of-bag (OOB) validation framework inherent to random forest algorithms, whereby bootstrap resampling is used to iteratively partition data into training and validation subsets across all trees within the model. This approach allowed all individuals to contribute to both model training and independent validation while avoiding the reduction in effective sample size associated with fixed train:test partitioning. Classification performance was assessed using overall OOB assignment accuracy, OOB error rates, and confusion matrices. Variable importance metrics were additionally examined to assess the relative contribution of δ^13^C and δ^18^O to classification success.

Unsupervised clustering analyses were then conducted to identify natural isotope assemblages independent of predefined waterway labels. Prior to analysis, isotope data were standardised to account for differences in variable scale. The optimal number of clusters was estimated using a gap statistic approach implemented in R with the factoextra package (Kassambara & Mundt 2026). Individuals were subsequently assigned to clusters using Partitioning Around Medoids (PAM) clustering implemented in the *cluster* package (Maechler *et al*. 2022).

Finally, uniform Manifold Approximation and Projection (UMAP) ordinations were was generated using the umap package (Konopka 2022) in R to visualise multivariate similarity among otolith isotope signatures in reduced two-dimensional space.

### 2.3 Mark-recapture and movement analysis

Between 2017 and 2020, a total of 868 mulloway were tagged across multiple estuarine and nearshore marine systems in south-eastern Australia as part of the citizen science–based mark–recapture program (Table 1). Tagging effort was concentrated in the Glenelg River, which accounted for approximately 63% of all tagged fish (n ≈ 550). Additional tagging occurred in the Patterson River (n ≈ 80), Hopkins River (n ≈ 40), Browns Bay (n ≈ 40), Yarra River (n ≈ 50), Coorong (n ≈ 25), Barwon River (n ≈ 20), Moyne River (n ≈ 13), and several smaller coastal and nearshore marine locations in western Victoria and south-eastern South Australia, including Rivoli Bay, Beachport, Kingston, and Port MacDonnell adjacent waters. Tagged individuals ranged from 38.5 to 140 cm total length (TL), with most fish between 50 and 70 cm TL.

Fish were tagged using standard external dart tags. Recapture records included recapture date, location, and total length at recapture, enabling estimation of movement distances, time at liberty, and growth. Minimum movement distance (km) was calculated as the river distance between release and the most recent recapture locations. A total of 172 tagged mulloway were recaptured during the study period, including 150 individuals recaptured once, 21 recaptured twice, and one individual recaptured four times. To maintain statistical independence, recapture histories were collapsed by fish ID such that only the most recent recapture event for each individual was retained for analyses. Individuals lacking complete movement, size, or time-at-liberty information were excluded from modelling, resulting in a final analytical dataset of 141 unique fish.

#### 2.3.1 Statistical analyses

To investigate factors associated with movement distance, movement analyses were conducted using Generalised Linear Models (GLMs) with a Gamma error distribution and log-link function implemented in the *stats* package in R version 4.2.1 (R Core Team 2022). This modelling approach was selected because movement distances were continuous, strictly positive, and highly right-skewed, with many individuals exhibiting short-distance movements and relatively few dispersing substantial distances. Predictor variables included fish total length (TL) and time at liberty (“days between capture and recapture”). Movement distributions were visualised using violin plots and scatterplots generated in R (v 4.2.1) using the ggplot2 (Wickham 2016) and tidyverse packages (Wickham *et al*. 2019). Model validation was conducted using residual diagnostics and dispersion assessment implemented in R (v 4.2.1) using the *DHARMa* (Hartig 2022) and *performance* (Lüdecke *et al*. 2021) packages. Statistical significance was assessed at α = 0.05.

### 2.4 Age structure and cohort composition analyses

Age estimates were obtained for 723 individuals from sagittal otoliths following established methods (Griffiths and Hecht, 1995; Silberschneider et al. 2009). Total length (mm) was also recorded for all individuals and used to examine spatial variation in age–length relationships among estuaries (Table S9). Briefly, one sagittal otolith from each fish was weighed (±0.001 g), embedded in resin, sectioned transversely through the core, and examined under transmitted light at 16× magnification. Annual growth increments (opaque zones) were counted along a transect extending from the primordium to the distal dorsal edge adjacent to the sulcus. Only complete marginal opaque zones followed by translucent material were included in counts. Age estimates were determined by Fish Ageing Services (FAS).

#### 2.4.1 Statistical analyses

Year of birth was estimated for each individual by subtracting estimated age from year of capture (Table S9). Reconstructed year-of-birth distributions were used as proxies for cohort strength and recruitment history. Cohort composition was visualised using annual histograms overlaid with kernel density estimates generated in ggplot2 (Wickham 2016), allowing identification of dominant year classes and comparison of recruitment patterns among estuaries and habitats. Estuarine cohort distributions were examined separately for waterways with sample sizes exceeding 20 individuals (Barwon, Glenelg, Hopkins and Patterson rivers; Table 1). Data manipulation and summary statistics were performed using functions from dplyr (Wickham et al. 2023). Differences in year-of-birth distributions among estuaries were assessed using Kruskal–Wallis rank-sum tests followed by Dunn’s post hoc comparisons with Benjamini–Hochberg correction implemented in the FSA package (Ogle et al. 2025). Differences in year-of-birth distributions and age structure between estuarine and marine habitats were evaluated using Wilcoxon rank-sum tests. Summary demographic statistics, including sample size, mean, median, standard deviation, and range, were calculated for each habitat to facilitate interpretation of habitat-specific age structure and recruitment patterns.

## 3 RESULTS

### 3.1 Genetic Analysis

A total of 314 mulloway individuals were successfully genotyped, with 2,010 SNP loci retained following filtering and used for downstream population genomic analyses (Table S1). Observed heterozygosity (*H*_O_) and expected heterozygosity (*H*_E_) were broadly comparable across all sampled waterways, indicating relatively similar levels of within-population genetic diversity throughout the study region (Table S1). Mean *H*_O_ across populations was 0.295 (range: 0.272–0.317), while mean *H*_E_ was 0.277 (range: 0.246–0.290). Estimates of *F*_IS_ were low across all populations (mean = −0.036; range = −0.106 to 0.016) and were predominantly slightly negative, indicating a small excess of heterozygotes relative to Hardy–Weinberg expectations (Table S1).

Global analyses revealed weak but significant population differentiation across the sampling range (*F*_ST_ = 0.043, 95% CI = 0.039–0.046), with pairwise *F*_ST_ estimates identified significant differentiation among locations from far west South Australia, south-east South Australia and central Victoria, and eastern Victoria and New South Wales, with little structure within these regional groupings (Figure 2a; Table S2). Consistent with these results, both DAPC and Bayesian clustering analyses identified three discrete genetic clusters corresponding to far west South Australia, south-east South Australia / central Victoria, and eastern Victoria / New South Wales (lowest BIC score = 3, Figure S1 and S2; Δ*K* = 3; Figure 2c). Hierarchical AMOVA further reinforced these concordant patterns, with 8.8% of genetic variation explained among these regions, compared with only 0.4% among populations within regions.

**Figure 2.**
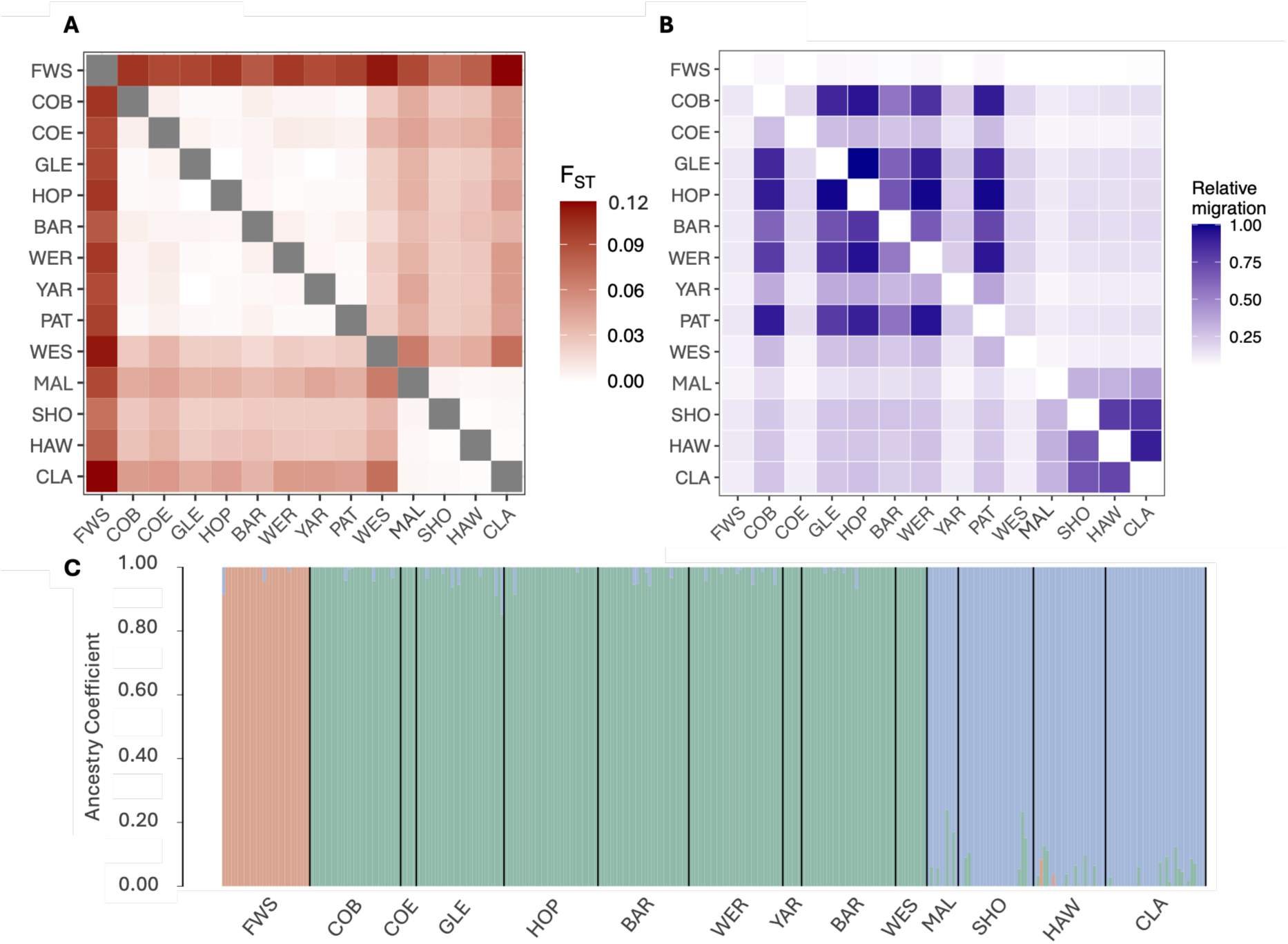
Estimates of population genetic structure and connectivity among mulloway populations across south-eastern Australia derived from complementary genomic approaches. (A) Heatmap of pairwise genetic differentiation (*F*_ST_), where darker shading indicates greater differentiation between population pairs. (B) Heatmap of relative migration rates inferred from genomic data, with rows representing source populations and columns recipient populations; darker shading indicates higher inferred migration and connectivity. Relative migration values are interpreted as low (<0.25), moderate (0.25–0.60), or high (>0.60). (C) Bayesian fineSTRUCTURE plot showing estimated ancestry membership coefficients for each individual across inferred population clusters, where ancestral clusters are defined by different colours. Each vertical bar represents an individual, partitioned according to proportional cluster membership. Individuals are grouped by sampling location and arranged geographically from west to east.

Relative migration analyses were broadly consistent with pairwise *F*_ST_ patterns and reinforced strong regional structuring across south-eastern Australia (Figure 2b; Table S3). Far West South Australia (FWS), which exhibited the highest pairwise *F*_ST_ values relative to eastern populations, also displayed uniformly weak inferred connectivity in both outgoing and incoming directions (range 0.04 - 0.13), supporting strong regional isolation and limited contemporary exchange. In contrast, the low pairwise *F*_ST_ values observed among Victorian estuaries were mirrored by moderate to strong inferred connectivity throughout the central south-eastern network, particularly among Coorong Beach, Glenelg River, Hopkins River, Barwon River, Werribee River, and Patterson River, where directional migration estimates frequently exceeded 0.8 (range 0.56 – 0.99), supporting stepping-stone connectivity and substantial shared genetic variation among adjacent estuaries. Connectivity estimates involving the Yarra River, Western Port Bay, and Coorong estuary were comparatively weaker (0.09–0.23) despite low pairwise *F*_ST_ values with neighbouring populations, suggesting that these waterways remain genetically similar to adjacent estuaries but may currently experience lower levels of directional exchange within the broader regional connectivity network. However, these patterns may also reflect the relatively small sample sizes for these locations and should therefore be interpreted cautiously. Strong inferred connectivity was also observed among Shoalhaven, Hawkesbury River, and Clarence River populations (range 0.68 – 0.90), consistent with low pairwise *F*_ST_ estimates and a broad eastern coastal connectivity corridor. In contrast, moderate estimates among Mallacoota and other east coast populations were observed (0.31–0.40) despite low pairwise *F*_ST_ estimates, suggesting more limited contemporary gene flow. Overall, directional connectivity within regions was generally characterised by strong bidirectional exchange rather than pronounced unidirectional gene flow, suggesting diffuse coastal connectivity and recurrent multi-directional dispersal among neighbouring estuaries. However, Hopkins River, Barwon River, Werribee River, and Patterson River consistently exhibited elevated outgoing connectivity toward adjacent systems, suggesting these waterways may occupy relatively central positions within the regional metapopulation network and potentially contribute disproportionately to regional recruitment.

### 3.2 Stable isotope analysis

#### 3.2.1 Overall spatial structuring among waterways

Otolith stable isotope measurements were successfully obtained from all 235 fish (Table S4). Using the full isotope dataset (n = 235), one-way ANOVA analyses identified highly significant differences among waterways for both δ^13^C (F = 38.90, p < 0.001) and δ^18^O (F = 11.86, p < 0.001), indicating substantial spatial heterogeneity in otolith isotope chemistry among estuarine systems across south-eastern Australia. Random forest supervised classification analyses achieved an overall out-of-bag (OOB) classification accuracy of 40.0%, corresponding to an OOB error rate of 60.0%, indicating relatively poor discrimination among waterways based on combined isotope signatures (Table 2). Variable importance analyses identified δ^13^C as contributing more strongly to classification performance than δ^18^O, accounting for 83.2% of total model importance relative to 16.8% for δ^18^O.

**Table 2.** Out-of-bag (OOB) confusion matrix from random forest supervised classification analyses of otolith stable isotope signatures (δ^13^C and δ^18^O) among mulloway capture waterways across south-eastern Australia. Rows represent the observed capture location of individuals, while columns represent the location predicted by the random forest classifier. Class error represents the proportion of individuals incorrectly assigned for each capture location.

|  | Coorong | Kingston | Glenelg River | Hopkins River | Barwon River | Yarra River | Westernport Bay | Patterson River | Gippsland Lakes | Mallacoota | Class.error |
| --- | --- | --- | --- | --- | --- | --- | --- | --- | --- | --- | --- |
| Coorong | 2 | 0 | 1 | 0 | 2 | 1 | 0 | 3 | 0 | 0 | 0.778 |
| Kingston | 0 | 0 | 0 | 0 | 2 | 1 | 0 | 0 | 0 | 0 | 1 |
| Glenelg River | 1 | 0 | 21 | 21 | 9 | 1 | 0 | 3 | 0 | 0 | 0.625 |
| Hopkins River | 1 | 0 | 22 | 28 | 3 | 1 | 0 | 0 | 0 | 0 | 0.491 |
| Barwon River | 0 | 1 | 6 | 2 | 28 | 3 | 0 | 13 | 0 | 1 | 0.481 |
| Yarra River | 0 | 0 | 2 | 1 | 2 | 1 | 0 | 3 | 0 | 0 | 0.889 |
| Westernport Bay | 0 | 0 | 0 | 0 | 0 | 0 | 3 | 2 | 0 | 0 | 0.4 |
| Patterson River | 3 | 0 | 4 | 1 | 13 | 3 | 1 | 11 | 1 | 2 | 0.718 |
| Gippsland Lakes | 0 | 0 | 0 | 0 | 0 | 0 | 0 | 2 | 0 | 0 | 1 |
| Mallacoota | 0 | 0 | 0 | 0 | 2 | 0 | 0 | 1 | 0 | 0 | 1 |

Unsupervised clustering analysis and the associated gap statistic identified two primary isotopic clusters within the dataset, representing the most parsimonious natural grouping structure in the isotope data (Figure S3). However, support for a more complex three-cluster solution was also evident (Figure S3). A UMAP ordination of the otolith isotope signatures revealed broad west–east spatial structuring among mulloway waterways across south-eastern Australia (Figure 3). Western waterways, including Coorong, Kingston, Glenelg River, and Hopkins River, formed a relatively distinct upper-left to central gradient within multivariate isotope space, indicating comparatively similar isotope profiles among these systems. In contrast, central Victorian waterways, including Barwon River, Yarra River, Patterson River, and Westernport Bay, occupied intermediate regions of the ordination and exhibited substantial overlap, suggesting weaker isotopic separation among neighbouring estuaries. Gippsland Lakes and Mallacoota individuals occupied more peripheral positions toward the upper-right region of the ordination, although some overlap with adjacent central Victorian assemblages remained evident.

**Figure 3.**
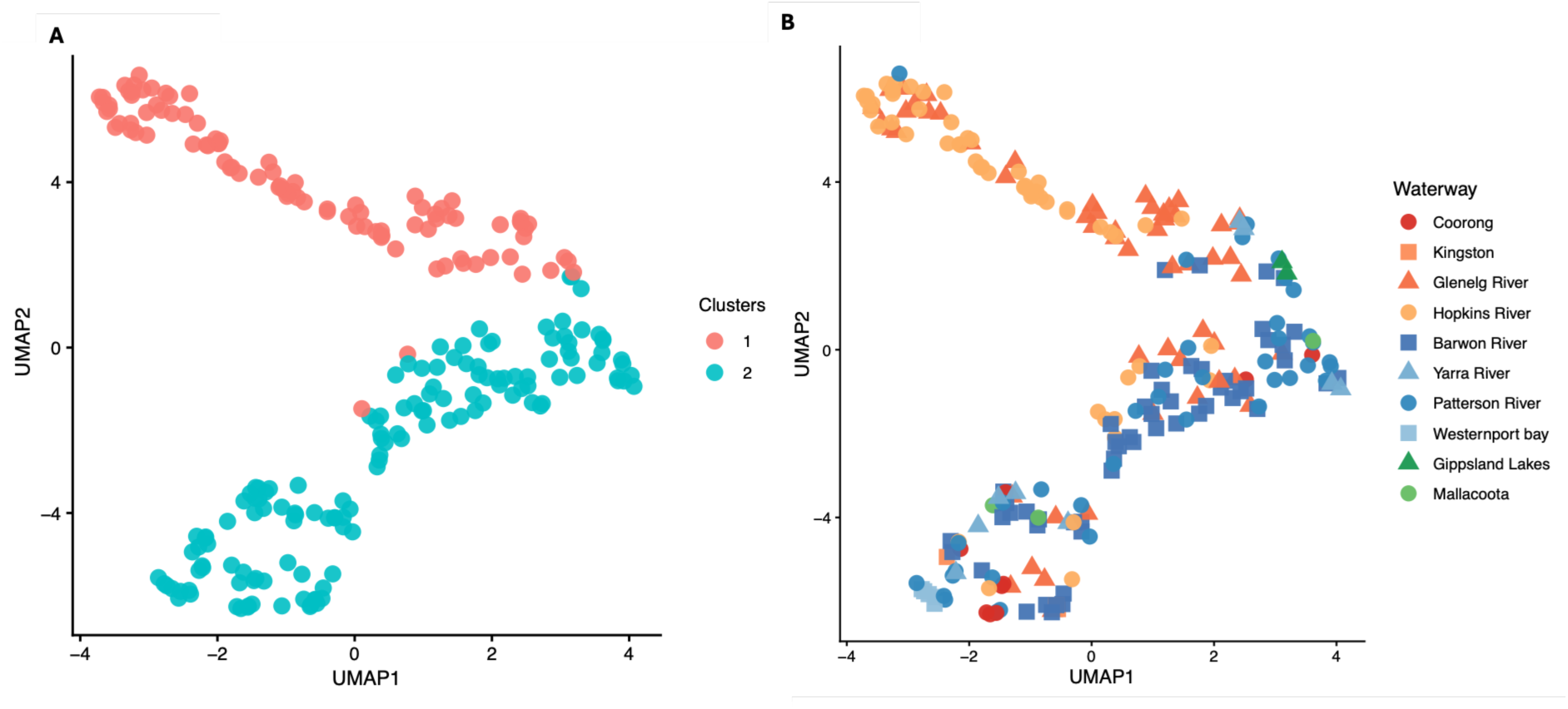
UMAP ordination of standardised otolith δ^13^C and δ^18^O isotope signatures for mulloway (*Argyrosomus japonicus*) across south-eastern Australia. (A) Individuals coloured according to the two clusters identified by partitioning around medoids (PAM) analysis. (B) The same ordination with individuals coloured and shaped according to capture waterway, illustrating overlap among central Victorian estuaries and greater differentiation among more geographically separated systems.

#### 3.2.2 Ontogenetic and sex-specific effects on isotope structuring

Exploratory ontogenetic analyses conducted on 149 individuals with complete age and total length data identified significant spatial variation in isotope chemistry among waterways for both sub-adults and adults. Sub-adults (n = 27) exhibited significant differences among waterways for both δ^13^C (F = 16.19, p < 0.001) and δ^18^O (F = 6.25, p = 0.007), while adults (n = 124) similarly displayed significant waterway differences for δ^13^C (F = 26.71, p < 0.001) and δ^18^O (F = 7.40, p < 0.001).

Subsequent supervised random forest classification analyses suggested stronger spatial discrimination among sub-adults relative to adults with sub-adults exhibiting comparatively high assignment accuracy (88.9%) and a low OOB error rate (11.1%; Table S5). In contrast, adults displayed substantially lower assignment accuracy (34.9%) and greater isotopic overlap among waterways within ordination space (Table S6). Variable importance analyses for sub-adults identified δ^13^C as contributing slightly more (55.4%) to classification performance than δ^18^O (44.6%). In contrast, adult classification models indicated substantially stronger influence of δ^13^C, which accounted for 87.6% of total model importance relative to 12.4% for δ^18^O.

Unsupervised clustering analyses similarly identified broad regional structuring among waterways within both ontogenetic stages. Gap statistic analyses identified a single cluster among sub-adults (Figure S4a), although UMAP ordinations still revealed some evidence for a separation between western and central Victorian waterways, with Glenelg River and Hopkins River individuals forming closely associated western clusters and Barwon River individuals occupying a broader and more distinct central Victorian region of ordination space (Figure S5a). In contrast, gap statistic analyses identified two primary isotopic clusters among adults (Figure S4b), with UMAP ordinations revealing broader overlap among western, central, and eastern waterways within multivariate isotope space (Figure S5b). Western-origin individuals from the Coorong, Kingston, Glenelg River, and Hopkins River occupied comparatively distinct regions of the ordination, while central Victorian waterways, including Barwon River, Yarra River, Patterson River, and Westernport Bay, formed broader overlapping assemblages. Eastern waterways, including Gippsland Lakes and Mallacoota, occupied more peripheral positions but still exhibited partial overlap with adjacent central Victorian assemblages. Overall, the greater isotopic overlap among adults suggests weaker spatial discrimination and increased regional mixing relative to sub-adults, although these ontogenetic differences should be interpreted cautiously given the substantial disparity in sample sizes between sub-adults (n = 25) and adults (n = 124).

Exploratory sex-specific analyses conducted on 123 individuals with available sex information similarly identified spatial structuring in isotope chemistry among waterways within both sexes. Female individuals (n = 75) exhibited significant waterway differences for both δ^13^C (F = 26.08, p < 0.001) and δ^18^O (F = 6.75, p < 0.001), whereas among males (n = 48) only δ^13^C varied significantly among waterways (F = 9.77, p < 0.001), with no significant differences detected for δ^18^O (F = 1.31, p = 0.273).

Supervised random forest classification analyses indicated stronger spatial discrimination among females relative to males, with females exhibiting higher assignment accuracy (63.2%; Table S7) than males (35.7%; Table S8). Variable importance analyses indicated relatively similar contributions of δ^13^C and δ^18^O to female classification performance (55.1% and 44.9%, respectively), whereas male classification models were driven primarily by δ^13^C, with δ^18^O exhibiting weak or negative importance values.

Unsupervised clustering analyses similarly identified broad regional structuring among waterways within both sexes, with gap statistic analyses identifying two primary isotopic clusters in both female and male datasets (Figure S6). UMAP ordinations revealed broad west–east spatial structuring among waterways in both sexes (Figure S7). Females exhibited comparatively stronger spatial separation, with western waterways, particularly Hopkins River and Glenelg River, forming a distinct cluster with limited overlap, whereas central Victorian waterways, including Barwon River, Patterson River, Yarra River, and Westernport Bay, occupied broader intermediate regions of ordination space. In contrast, males displayed greater isotopic overlap among waterways, although Glenelg River and Hopkins River still formed a comparatively distinct western cluster separated from the broader central Victorian gradient. Coorong, Gippsland Lakes, and Mallacoota individuals occupied more peripheral positions in both sexes but were represented by comparatively few samples. Overall, the ordinations suggest weaker spatial discrimination and greater isotopic overlap among males relative to females. However, these apparent sex-specific differences should be interpreted cautiously given unequal sample sizes between females (n = 75) and males (n = 48), which may influence clustering structure and assignment success.

### 3.3 Mark-recapture and movement analysis

Approximately 70.2% of recaptured mulloway (99 of 141 fish) were ultimately detected within the same river or estuarine system in which they were originally tagged, demonstrating strong estuarine fidelity and local residency (Table 3; Table S9). Within-estuary recaptures were recorded across the Glenelg (n = 90), Patterson (n = 18), Hopkins (n = 5), Moyne (n = 2), and Yarra rivers (n = 5), with fish remaining within the same system for 48–755 days between tagging and recapture (Table S9). Overall, 76 fish (53.9%) were ultimately recaptured within 5 km of their original tagging location, while a further 34 fish (24.1%) were recaptured >5 km from their tagging location but still within the same river or estuarine system, supporting prolonged use of local habitats and relatively restricted home ranges following recruitment (Figure 4). When all recapture events were considered (including repeated recaptures of the same individuals), evidence for local residency was even stronger, with 94 of 151 recapture events (62.3%) occurring within 5 km of the original tagging location and a further 35 events (23.2%) involving movements >5 km but remaining within the original river or estuarine system (Figure 4).

**Figure 4.**
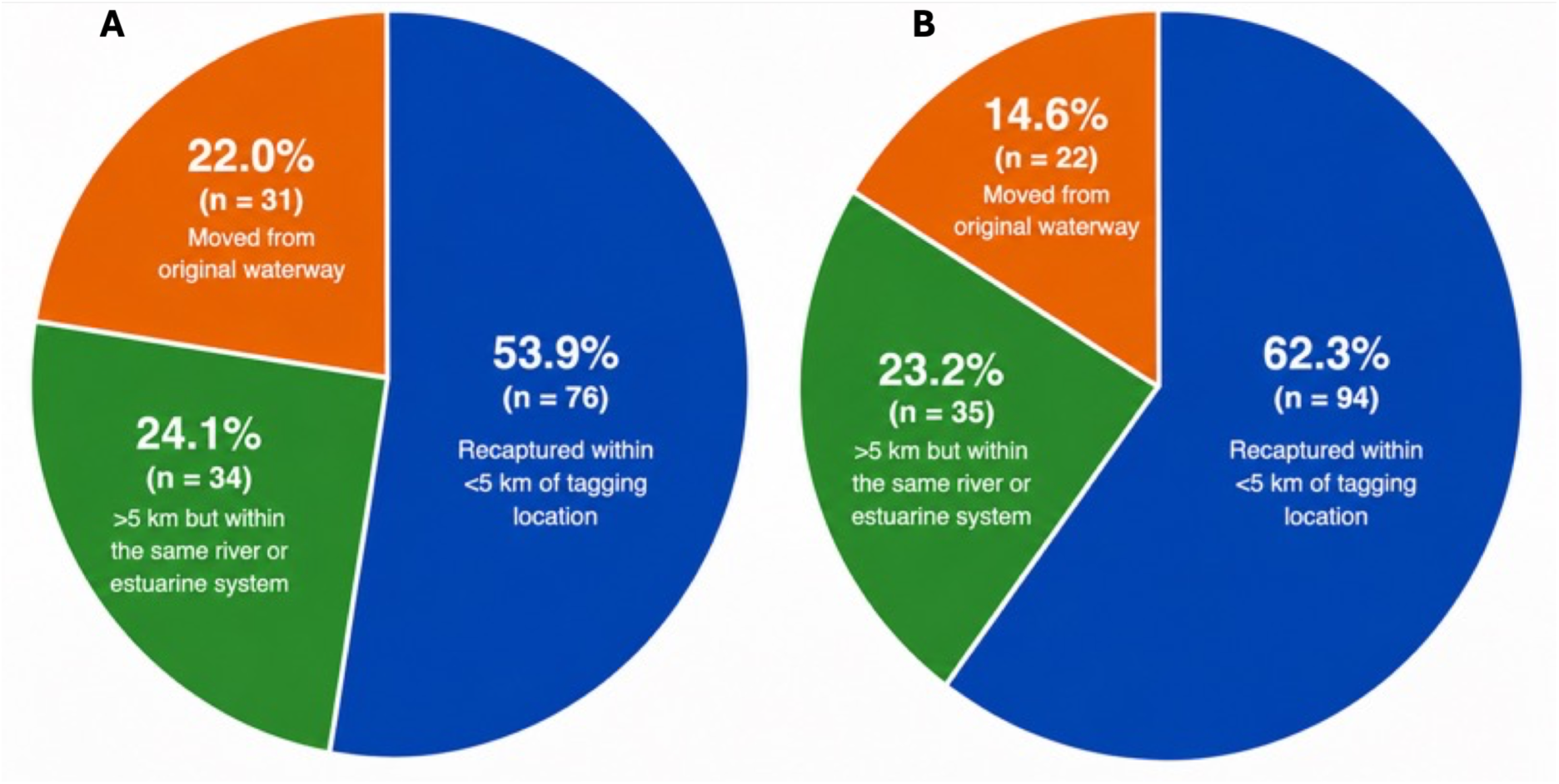
Recapture outcomes of tagged mulloway (*Argyrosomus japonicus*) based on (A) final recapture location for each individual fish (n = 141), and (B) all recapture events, including repeated recaptures of the same individuals (n = 294 recapture events).

**Table 3.** Summary of mulloway (*Argyrosomus japonicus*) recapture movement pathways based on all recorded recapture events and final recapture locations.

| Movement type | Specific movement | Recapture |  |
| --- | --- | --- | --- |
|  |  | Sub-Total | Total |
| 1. Within estuary | Glenelg River | 90 | 120 |
|  | Hopkins River | 5 |  |
|  | Moyne River | 2 |  |
|  | Patterson River | 18 |  |
|  | Yarra River | 5 |  |
| 2. Between Estuaries | Patterson River → Yarra River | 1 | 3 |
|  | Yarra River → Maribyrnong River | 1 |  |
|  | Glenelg River → Coorong Lagoon | 1 |  |
| 3. Estuary to Coastal Waters | Glenelg River → Port MacDonnell (SA) | 3 | 8 |
|  | Glenelg River → Ocean Grove | 1 |  |
|  | Moyne River → Port Fairy | 1 |  |
|  | Barwon River → Coorong beach | 1 |  |
|  | Patterson River → Port Phillip Bay | 2 |  |
| 4. Coastal Waters to Estuary | Port MacDonnell, SA → Glenelg River | 11 | 18 |
|  | Coorong beach, SA → Glenelg River | 1 |  |
|  | Coorong beach, SA → Coorong Lagoon | 4 |  |
|  | Barwon Coast → Barwon River | 2 |  |
| 5. Coastal Waters | Port MacDonnell → Port MacDonnell | 1 | 2 |
|  | Port MacDonnell → Coorong beach | 1 |  |

Despite this predominance of local retention, a substantial proportion of individuals exhibited long-distance movements, highlighting episodic connectivity among waterways. Approximately 22.0% of recaptured mulloway (31 of 141 fish) were recaptured outside their original capture waterway (Figure 4). Across all recapture events 14.6% (22 of 151) involved movements away from tagging location (Figure 4), indicating that while most individuals exhibited strong estuarine fidelity, a smaller more mobile subset disproportionately contributed to broader-scale connectivity among estuarine and coastal systems. These inter-waterway movements occurred predominantly among fish between 50 and 80 cm total length (TL), including nine individuals within the 50–59 cm size class, 15 within the 60–69 cm size class, seven within the 70–79 cm size class, and seven within the 80–89 cm size class. Fewer dispersers were recorded among smaller (40–49 cm; *n* = 1) and larger fish (90–99 cm; *n* = 2). Movement pathways included transfers among neighbouring western Victorian estuaries, movements between estuaries and adjacent coastal marine waters, and broader-scale coastal dispersal between Victoria and south-eastern South Australia (Table 3). Based on final recapture locations, inter-system movements included three transfers between estuaries, eight movements from estuaries into coastal waters, and 18 movements from coastal waters into estuarine systems, demonstrating strong bidirectional connectivity between estuarine and marine habitats. Notable examples included movements from the Glenelg River to the Coorong Lagoon (∼420 km) and the largest recorded displacement from the Barwon River to the Coorong (∼700 km).

Gamma GLM analyses identified weak positive relationships between total movement distance, fish body size, and time at liberty. Larger fish tended to move greater distances than smaller individuals (β = 0.060 ± 0.032 SE, z = 1.87, P = 0.061; Figure 5), while fish that remained at liberty for longer periods also tended to exhibit greater displacement distances (β = 0.0032 ± 0.0018 SE, z = 1.81, P = 0.071; Figure 5). Although neither predictor was statistically significant at α = 0.05, both relationships showed consistent positive trends. The overall explanatory power of the model was low (Pseudo-*R*² = 0.005), indicating substantial unexplained variation among individuals. Residual diagnostics indicated acceptable model performance, with no evidence of substantial overdispersion (dispersion ratio = 1.02) or major deviations from model assumptions based on simulated residual diagnostics.

**Figure 5.**
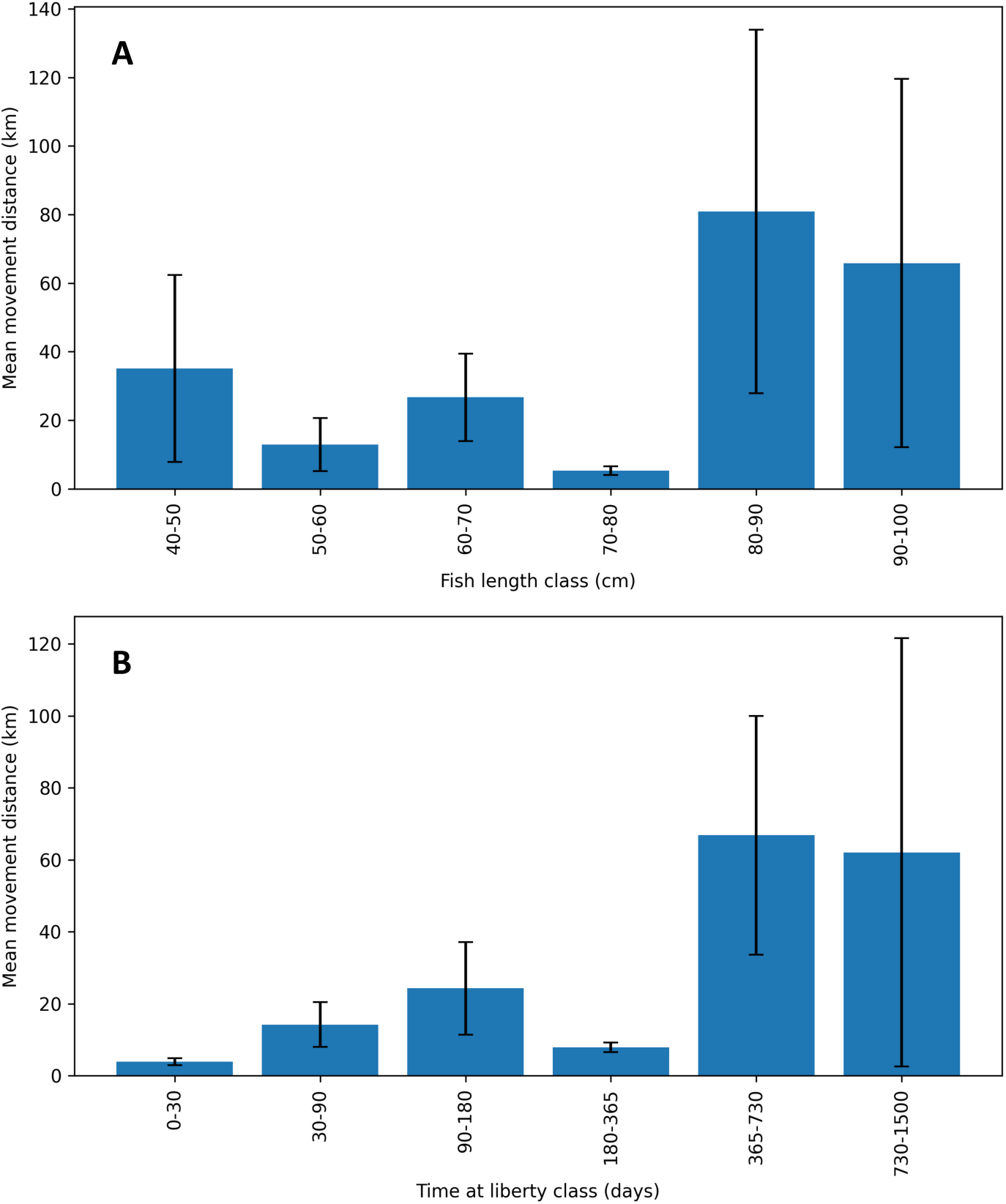
Mean total movement distance of recaptured mulloway (*Argyrosomus japonicus*) across (A) fish length classes (10 cm bins) and (B) time-at-liberty classes. Bars represent mean movement distance (km) ± SE based on the displacement between original tagging location and most recent recapture location for each individual fish. Analyses included only unique recaptured individuals, with repeated recaptures collapsed to the most recent observation per fish (n = 141).

### 3.4 Age structure, cohort composition and growth analyses

Year-of-birth distributions of estuarine fish were strongly concentrated between 2009 and 2013, with a pronounced peak centred on 2011–2012, indicating dominance by a relatively narrow range of recent cohorts (Figure 6A). Only a small number of individuals were assigned to cohorts prior to 2005. Cohort distributions were broadly similar among estuaries, with all four major estuaries exhibiting dominant cohorts between 2010 and 2013. Although year-of-birth distributions differed overall among estuaries (Kruskal–Wallis χ² = 11.73, df = 3, P = 0.008), no pairwise comparisons remained significant following correction for multiple testing (all adjusted P > 0.05), suggesting that recruitment histories were largely synchronous a cross systems and that differences in cohort composition were subtle.

**Figure 6.**
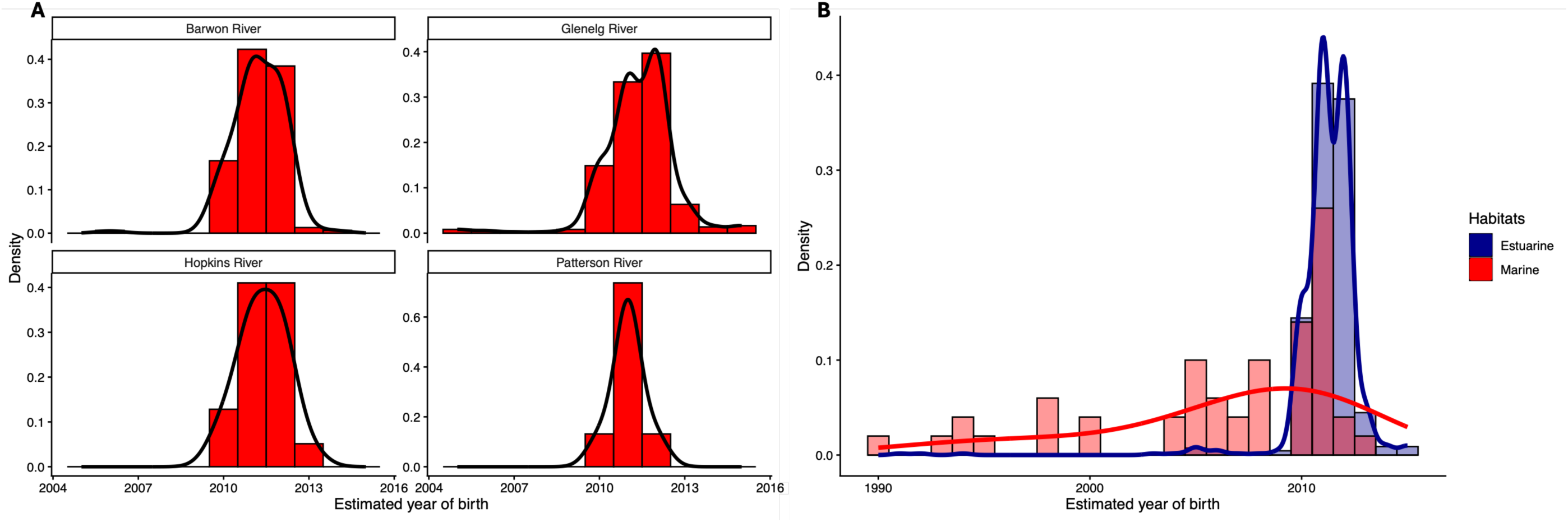
Cohort composition and age structure of mulloway (*Argyrosomus japonicus*) sampled from south-eastern Australia. (A) Year-of-birth distributions for estuarine fish from the Barwon River, Glenelg River, Hopkins River, and Patterson River. (B) Comparison of year-of-birth distributions between estuarine (n = 672) and marine (n = 50) fish. Histograms depict annual cohort frequencies and solid lines represent kernel density estimates.

Marked differences in cohort composition and age structure were evident between habitats (Figure 6B). Estuarine fish (n = 672) were dominated by recent cohorts, with a mean year of birth of 2011, whereas marine fish (n = 50) exhibited a broader cohort distribution extending from 1990 to 2013, with a mean year of birth of 2008. Year-of-birth distributions differed significantly between habitats (Wilcoxon rank-sum test, W = 28,072, P < 0.001). Estuarine fish were also younger on average (5.2 ± 1.4 years; median = 5 years) than marine fish (9.1 ± 5.9 years; median = 6 years), and age distributions differed significantly between habitats (Wilcoxon rank-sum test, W = 8,133.5, P < 0.001).

## 4 DISCUSSION

This study integrates genomics, otolith chemistry, and mark–recapture analyses to resolve connectivity in mulloway across ecological and evolutionary timescales. Populations across south-eastern Australia were characterised by strong local estuarine residency alongside broader regional connectivity maintained through intermittent dispersal and episodic recruitment events. Genomic analyses revealed sufficient gene flow among waterways to limit population genetic divergence across much of the study area, whereas otolith isotope chemistry identified substantial environmental heterogeneity and fine-scale ecological structuring among estuaries, consistent with sustained occupancy of local estuarine habitats. Mark–recapture analyses supported this interpretation, demonstrating high estuarine fidelity in most fish despite occasional long-distance coastal movements by a smaller subset of individuals. Finally, reconstructed age structures revealed a shared cohort signature across estuaries, with most fish originating from a single strong recruitment pulse centred on 2010–2011. Together, our findings indicate that mulloway fisheries comprise regionally connected networks of partially independent estuarine stocks, where strong local residency is intermittently offset by dispersal and episodic recruitment that maintain long-term demographic and genetic connectivity. Despite this broader connectivity, strong estuarine fidelity suggests local populations, at least within Victorian estuaries, may be vulnerable to localised depletion under sustained fishing pressure and reductions in freshwater flows. These patterns highlight the need for conservative, spatially explicit management that protects estuarine habitats, maintains connectivity pathways, and reduces the risk of local depletion. More broadly, the study demonstrates how integrating complementary approaches provides a stronger framework for resolving population structure and identifying biologically meaningful management units in exploited fisheries with complex life histories.

### 4.1 Complex stock connections

Our results reveal a disconnect between intergenerational gene flow and contemporary demographic exchange in mulloway across south-eastern Australia. Population genomic analyses indicated that little genetic differentiation among mulloway stocks across more than 900 km of coastline spanning south-east South Australia and Victoria. Consistent with Barnes et al. (2016), these findings suggest substantial long-term connectivity likely maintained through dispersal during early life stages. However, significant genetic differentiation was evident between regions influenced by different current systems, with stocks from far west South Australia, south-east South Australia/Victoria, and New South Wales comprising isolated populations. The connectivity break between far western and central regions likely reflects the large geographic distances between sites and the paucity of estuarine habitats along this coastline. In contrast, the break between central and New South Wales populations aligns with the well-recognised biogeographic divide in eastern Victoria associated with Bass Strait and the historical Bassian Isthmus (York *et al*. 2008; Colgan & Schreiter 2011; Miller *et al*. 2013).

In contrast to the relatively weak genomic structure, significant differences in both δ^13^C and δ^18^O among mulloway from separate waterways reflect contrasting environmental and/or dietary histories among estuaries. Supervised random forest classification analyses achieved moderate to high assignment success, indicating that many individuals retained isotope signatures characteristic of their capture waterway. Unsupervised clustering analyses additionally identified two primary isotopic assemblages, broadly separating South Australian and Victorian waterways from those on Australia’s east coast. Central Victorian estuaries, however, exhibited substantial overlap in isotope space, potentially reflecting shared environmental conditions, hydrological similarity, or greater exchange among adjacent systems. Stronger spatial discrimination among sub-adults relative to adults further suggests that younger fish occupy more localised estuarine habitats with unique thermal regimes and basal carbon types supporting the food-web or metabolism (Campana 2005; Elsdon *et al*. 2008). In contrast, adults likely integrate isotope signals across broader spatial scales through increased movement. Despite reduced statistical power arising from uneven sample sizes among life stages, these findings are consistent with previous otolith microchemistry studies indicating ontogenetic shifts in residency patterns, with juveniles exhibiting greater estuarine fidelity and adults becoming increasingly dispersive (Taylor *et al*. 2006a; Silberschneider & Gray 2008; Hughes *et al*. 2022). Preliminary sex-specific analyses also suggested somewhat stronger spatial structuring among females than males, although unequal sample sizes limited strong inference. Overall, our results corroborate previous otolith-based studies demonstrating distinct estuarine and regional chemical signatures in mulloway, moderate assignment success to nursery or regional origin, and evidence of regional stock structuring across eastern and southern Australia (Ferguson *et al*. 2011; Russell *et al*. 2021). Collectively, these findings support prolonged estuarine residency alongside partial regional connectivity, with connectivity dynamics varying across ontogeny and potentially among sexes.

Differences between genomic and isotope patterns highlight the contrasting temporal and ecological resolution provided by each method which are well recognised in the literature (Cadrin *et al*. 2013; Sarakinis *et al*. 2024). Population genomic connectivity estimates primarily reflect realised gene flow integrated across multiple generations, whereas otolith isotope chemistry captures more contemporary environmental exposure and habitat occupancy at the scale of individual fish. Consequently, weak genomic differentiation does not necessarily imply frequent contemporary movement among estuaries. Instead, our combined results suggest that many mulloway exhibit prolonged local residency sufficient to develop estuary-specific isotope signatures, while occasional dispersal and recruitment events remaining adequate to maintain broader regional connectivity and genetic homogeneity. The apparent discrepancy between weak genomic structure and stronger isotope differentiation therefore reflects the complementary rather than conflicting nature of these approaches.

Mark–recapture analyses provided direct behavioural evidence supporting our interpretation of mulloway stock structure. Approximately 70% of recaptured fish were recorded within their original estuarine systems, and over half were recaptured within 5 km of their tagging location, demonstrating strong estuarine fidelity and relatively restricted movement for many individuals. Some fish were recaptured in the same estuary more than two years after tagging, indicating prolonged use of local habitats following recruitment. While these observations cannot exclude temporary movements beyond the capture system between release and recapture, they nonetheless suggest that many individuals spend extended periods associated with particular estuaries. These findings reinforce the isotope results by demonstrating that environmentally distinct isotope signatures can develop and persist through prolonged occupancy of localised estuarine habitats. Strong local residency also suggests that estuarine systems function as important long-term habitats rather than merely transient nursery areas. These patterns closely align with previous large-scale tagging studies of eastern Australian mulloway, which similarly reported that most individuals were recaptured within their estuary of release or within short coastal distances (Hughes *et al*. 2022). However, movement behaviour may be context dependent. For example, Barnes *et al*. (2019) reported that mulloway in South Australia’s far west coastal region remained close to tagging locations during summer but undertook migrations of up to 550 km in autumn. Together, these findings suggest that although long-distance movements can occur and may be seasonally important, many mulloway exhibit strong site fidelity and prolonged associations with particular estuarine habitats.

Despite the predominance of local residency, mark–recapture data identified substantial long-distance dispersal by a smaller subset of individuals. Approximately one-fifth of recaptured fish moved outside their original estuarine systems, including several coastal movements spanning hundreds of kilometres. Movements occurred both among neighbouring estuaries and between estuarine and coastal marine habitats, demonstrating bidirectional connectivity between these environments. Particularly notable were movements between Victorian estuaries and the Coorong region of South Australia, including one displacement approaching 700 km. These patterns are consistent with previous cooperative tagging studies reporting predominantly localised movements punctuated by infrequent but ecologically important coastal migrations exceeding 500 km, including movements between the Glenelg River and Coorong estuary (Barnes *et al*. 2019; Lieschke 2019; Hughes *et al*. 2022). Unlike previous studies suggesting strong ontogenetic influences on movement (Taylor *et al*. 2006b; Ferguson *et al*. 2011; Russell *et al*. 2021; Hughes *et al*. 2022), body size did not correspond to routine inter-estuary dispersal. Although larger mulloway generally exhibited greater movement distances, the relationship was weak and highly variable, with substantial overlap among size classes. Most adults were repeatedly recaptured within their original systems, indicating prolonged association with particular estuarine or coastal regions, while broader coastal and inter-system dispersal was undertaken by only a minority of more mobile individuals. Together, these rare but consequential dispersal events likely contribute to the relatively weak genomic differentiation observed over large spatial scales.

Overall, our findings closely mirror those reported for mulloway (*A. japonicus*) populations in South Africa. Genetic studies there similarly identified weak differentiation across more than 2000 km of coastline, despite strong evidence of estuarine fidelity and restricted movement at finer spatial scales (Mirimin *et al*. 2016). Behavioural studies further demonstrated high juvenile residency within estuarine nursery habitats and little evidence of persistent longshore dispersal or annual spawning migrations (Næsje *et al*. 2012, Cowley *et al*. 2013; Dunlop *et al*. 2013). More recently, Knight *et al*. (2025) analysing nearly four decades of tag–recapture data, reported predominantly localised movements among juveniles and sub-adults, with only a small proportion of adults undertaking larger-scale movements consistent with spawning migrations. Together, these studies support an emerging paradigm of partial migration and metapopulation structure in *A. japonicus*, where broad-scale genetic connectivity is maintained by occasional dispersive individuals and episodic recruitment events, while strong local residency generates demographic independence among estuaries and coastal regions.

### 4.2 Evidence of recruitment constraits

Reconstructed year-of-birth distributions revealed remarkably similar cohort composition among estuaries, with all major systems exhibiting a pronounced recruitment peak centred on 2011–2012, indicating synchronous recruitment histories across Victorian estuaries. This synchrony suggests that recruitment dynamics are driven by regional-scale processes rather than independent local events. Notably, these dominant cohorts coincided with the major Murray River floods of 2010–2011, which have previously been linked to mass mulloway spawning and recruitment in the Coorong, with subsequent populations largely comprising the direct progeny of spawning associated with this flood event (Bice et al. 2012). Independent evidence from New South Wales similarly indicates that increased rainfall and river flows from 2008 onwards enhanced mulloway recruitment across multiple estuaries, highlighting the importance of broad-scale hydrological conditions during the spawning period (NSW Department of Primary Industries 2016). Together, these findings support the hypothesis that episodic high-flow events generate regionally strong recruitment pulses that are dispersed among estuarine nursery habitats, contributing to demographic connectivity and the absence of detectable genetic structure across most of southern and eastern Australia.

Estuarine and marine habitats exhibited markedly different demographic structures, with estuaries dominated by younger fish and recent cohorts, while marine environments contained a broader range of cohorts and significantly older individuals. This pattern is consistent with the recognised role of estuaries as critical nursery habitats for mulloway and provides strong evidence for ontogenetic shifts from estuarine to coastal marine environments (Taylor et al. 2006a; Silberschneider & Gray 2008; Hughes et al. 2022). Across eastern Australia, juvenile mulloway are strongly associated with estuarine habitats where growth rates are high and predation risk is reduced, often remaining within these systems for several years before progressively moving into coastal marine environments as they approach maturity (Gray & McDonall 1993; Taylor *et al*. 2006a; Silberschneider & Gray 2008). Similar patterns have been documented in South Africa, where conspecifics (*A. japonicus*) are concentrated in estuaries and surf-zone nursery habitats, while larger adults predominantly occupy coastal and offshore environments and undertake seasonal spawning aggregations (Griffiths 1996; Cowley *et al*. 2008). The predominance of younger cohorts within estuaries and older age classes in marine habitats observed here therefore reflects a broadly conserved life-history strategy throughout the species’ range, whereby estuaries function as recruitment and juvenile development areas before individuals transition to marine habitats for maturation and reproduction. This ontogenetic habitat shift likely contributes to the demographic differences observed between environments and reinforces the importance of estuarine nursery habitats in sustaining regional mulloway populations.

### 4.2 Implications for fisheries management

Together, our results suggest that mulloway populations from south-eastern South Australia and Victoria function as a partially connected metapopulation, with most individuals contributing to largely discrete estuarine stocks. These stocks therefore appear neither fully independent nor part of a single homogeneous regional population, but instead comprise semi-discrete demographic units linked by intermittent dispersal, and episodic spawning and irregular recruitment. The strong local residency observed in Victorian stocks suggests depleted estuarine populations may not be rapidly replenished by immigration, particularly where dispersal is infrequent and driven by relatively few individuals, and recruitment is irregular and flow dependent. The consistency of our findings with studies from other regions suggests that these dynamics are characteristic of mulloway populations throughout southern and eastern Australia, with the notable exception of the far west South Australian population, which exhibits a distinct life-history strategy characterised by limited estuarine association (Ferguson 2010; Barnes et al. 2016). For fisheries management, this highlights the risk of local depletion and underscores the need for spatially explicit strategies that protect local stocks, maintain estuarine habitat quality and nursery areas, and preserve coastal connectivity pathways that support demographic exchange and long-term genetic resilience.

As previously discussed, infrequent, flow-driven spawning and recruitment events likely play a disproportionate role in sustaining regional connectivity, genetic diversity, and replenishment of estuarine mulloway stocks. Altered freshwater flow regimes have long been previously recognised as a key threat to the sustainability of mulloway fisheries, with river regulation, dam construction, water extraction, and reduced flood frequency likely to weaken or disrupt the environmental cues associated with spawning and recruitment (Ferguson *et al*. 2008; Taylor *et al*. 2014). Reduced freshwater inflows may also compress estuarine salinity gradients and reduce the extent of brackish nursery habitats relied upon by juveniles (Gray & Mcdonall 1993; Ferguson *et al*. 2008; Taylor *et al*. 2014). These pressures are likely to intensify under climate change, with projections for south-eastern Australia indicating declines in freshwater discharge, longer droughts, reduced runoff, and fewer moderate flood events despite occasional increases in extreme rainfall intensity (Gillanders *et al*. 2022; CSIRO and Bureau of Meteorology 2022). Such changes may reduce the frequency and predictability of recruitment-triggering flow events, weakening connectivity and stock replenishment among estuaries and increasing the risk of localised or serial depletion.

These concerns are particularly relevant given the historical depletion and ongoing rebuilding of mulloway stocks in parts of south-eastern Australia, especially in New South Wales, where recovery has relied heavily on conservative harvest controls and sporadic strong recruitment years (NSW Department of Primary Industries 2016; DPIRD 2025). More broadly, increasing recreational fishing pressure, estuarine habitat degradation, and climate- driven reductions in freshwater inflow may further limit the resilience of local populations by reducing both spawning output and the frequency of successful recruitment events. Management strategies should therefore account for recruitment variability and environmental stochasticity through precautionary harvest controls, protection of flow-dependent spawning processes, and adaptive management frameworks (e.g. genetically informed stock augmentation; (Weeks *et al*. 2011; Hoffmann *et al*. 2021) capable of responding to fluctuating recruitment and connectivity dynamics through time.

### 4.3 Integrating Connectivity Metrics for Effective Fisheries Management

The integration of complementary methods represents a major strength of this study and provides a more complete understanding of fisheries connectivity than any single approach alone. Genomic analyses quantified long-term realised gene flow, isotope chemistry characterised contemporary habitat occupancy and estuarine environmental structure, and mark–recapture analyses directly measured individual movement behaviour, together revealing a nuanced connectivity framework in which ecological residency and evolutionary connectivity coexist simultaneously. Genomics alone may have underestimated the ecological importance of local estuarine fidelity, whereas movement data alone may have overstated demographic isolation by failing to capture the long-term significance of infrequent dispersal. More broadly, our findings highlight the importance of integrated, multidisciplinary approaches for assessing connectivity in fishes with complex life histories, particularly where low-frequency dispersal maintains genetic homogeneity while post-recruitment behaviour generates fine-scale demographic structure (Cadrin *et al*. 2013). In such systems, reliance on genetic data alone may overestimate functional connectivity and resilience, potentially masking the risk of localised depletion, whereas movement or habitat- use studies in isolation may underestimate the replenishment capacity provided by rare dispersal events. Integrating intergenerational and contemporary connectivity metrics therefore provides a more realistic assessment of stock structure, resilience, and risk, and offers a stronger foundation for spatially explicit fisheries management.

## Acknowledgements

This project was funded by the Victorian Fisheries Authority through its Large and Small Recreational Fishing Grants Programs. We sincerely thank the many recreational anglers who contributed samples and recapture data through the citizen science program, without whom this research would not have been possible. We are also grateful to fisheries officers Matt Proctor, Rick Moreton, and Trudy Schmidt for their assistance with sample collection, logistics, and storage, and to Matt Taylor and Julian Hughes for assistance with sample collection and processing. We thank the staff at Fish Ageing Services for their support, expertise, and for providing access to their facilities to undertake ageing of mulloway specimens.

## Author contributions

This project was conceived by L.B., A.D.M., and N.S.W. The citizen science mark–recapture study and data acquisition were led by L.B. and N.S.W., while tissue collections were led by L.B. and T.C.B. Data analyses were led by A.D.M. with assistance from L.B., J.R., and J.M. A.D.M. led manuscript preparation with contributions from all authors.

## Data Availability Statement

All genomic datasets are publicly available in the DRYAD archives

